# A novel long-amplicon *rpoB* primer pair for high resolution microbiome analysis at the species-level

**DOI:** 10.64898/2026.05.15.725465

**Authors:** Marc Venbrux, Sam Crauwels, Hans Rediers

**Author notes:** Address correspondence to Hans Rediers.

## Abstract

The 16S rRNA gene is the most widely used genetic marker for microbial community profiling, but its limited sequence divergence often prevents species-level identification. The RNA polymerase β-subunit gene (*rpoB*) offers higher sequence variability, single-copy occurrence, and stronger phylogenetic consistency, yet its adoption in metataxonomic studies has been constrained by the lack of universal primer sets. Here, we present a novel universal primer pair that amplifies an ∼1,800 bp *rpoB* region (rpoB_MV) compatible with long-read sequencing platforms. *In silico* evaluation across 17683 bacterial reference genomes demonstrated high universality, with over 86% of genomes predicted to amplify. Compared with full-length and partial 16S rRNA gene markers, the rpoB_MV amplicon exhibited significantly greater inter-species sequence divergence and improved phylogenetic concordance with core-genome trees. Sequencing of two complementary mock communities confirmed superior species-level identification accuracy, with misclassification rates below 0.01% and no reads assigned to unresolved species clusters. These results establish rpoB_MV as a robust alternative to 16S rRNA gene-based profiling for high-resolution metataxonomic applications.

**IMPORTANCE:** Microbial community studies increasingly require species-level resolution because species within the same genus can differ substantially in pathogenicity, ecological function, and metabolic capacity. Current 16S rRNA gene-based methods frequently fail to distinguish closely related species, collapsing biologically distinct organisms into the same taxonomic assignment and obscuring community differences that matter for clinical diagnostics, food safety, and environmental monitoring. The rpoB_MV primer pair presented here overcomes this limitation by targeting a longer, more variable region of the *rpoB* gene, enabling accurate species-level identification across diverse bacterial phyla. Combined with advances in long-read sequencing, this approach provides researchers with a practical tool to resolve microbial communities at the species-level.

## INTRODUCTION

Metataxonomics, also known as metabarcoding, is widely used to study microbial communities by amplifying and sequencing conserved genetic markers from environmental DNA (1). For bacteria, the most commonly used genetic marker is the 16S rRNA gene, which contains nine hypervariable regions (V1-V9) (2). However, short-read sequencing generally captures only one or two hypervariable regions, and the choice of region can influence taxonomic performance across bacterial taxa (2–5). Long-read platforms such as PacBio and Oxford Nanopore sequencing now enable high-throughput sequencing of the nearly full-length 16S rRNA gene (∼1450 bp), improving resolution compared with partial 16S rRNA gene sequencing (2, 6). Nevertheless, the 16S rRNA gene has important limitations that affect the accuracy and resolution of microbial community analyses. Its sequence variation is often insufficient to distinguish closely related species, such as *Bacillus cereus* and *Bacillus anthracis* (7, 8). This hampers accurate interpretation of community composition when closely related species differ in biological function or virulence. In addition, variable 16S rRNA gene copy number and intragenomic sequence variability can complicate interpretation of sequencing data and bias estimates of relative abundance and species richness (8).

As an alternative to the 16S rRNA gene, conserved housekeeping genes can overcome some of these limitations because they are typically single-copy and exhibit higher sequence variability. One promising marker is *rpoB*, which encodes the RNA polymerase β-subunit, has been shown to provide higher taxonomic resolution than partial or full-length 16S rRNA gene sequences (8–10). A disadvantage of *rpoB*, and protein-encoding genes in general, is that universal primer design is difficult due to silent third codon mutations (10). Few universal primer sets exist for *rpoB*, and those available are often restricted to particular taxonomic groups or amplify short fragments that do not exploit long-read sequencing (11–13). For example, recently described universal *rpoB* primer pairs amplify 434- and 535-bp regions and were developed for Illumina or Sanger sequencing workflows (11, 12).

To the best of our knowledge, no broadly validated universal primers are currently available for amplification and long-read sequencing of large bacterial *rpoB* regions. Here, we designed and validated a universal primer pair targeting an approximately 1.8 kb *rpoB* region for species-level metataxonomic profiling. We assessed primer universality *in silico* and experimentally, benchmarked this novel genetic marker against the 16S rRNA gene and previously described *rpoB* markers, and evaluated its performance using nanopore sequencing of two mock communities.

## METHODS

### Bacterial strains and mock communities used in this study

A representative panel of 20 bacterial strains was selected for PCR validation and construction of an equimolar mock community (Mock20; Table 1). The panel, selected to represent diverse phylogenetic groups, covered five bacterial phyla, including *Pseudomonadota*, *Bacillota*, *Bacteroidota*, *Actinomycetota*, and *Campylobacterota*, and included two *Xanthomonas* species to evaluate discrimination of closely related taxa. Strains were cultured under species-appropriate conditions, and genomic DNA was extracted using the DNeasy PowerLyzer Microbial Kit (Qiagen). DNA concentration was determined using a Qubit 4 fluorometer with the dsDNA High Sensitivity Assay Kit (Thermo Fisher Scientific). Genomic DNA for *Campylobacter coli* and *Helicobacter pylori* was obtained directly from the Belgian Coordinated Collections of Microorganisms (BCCM).

**Table 1.**
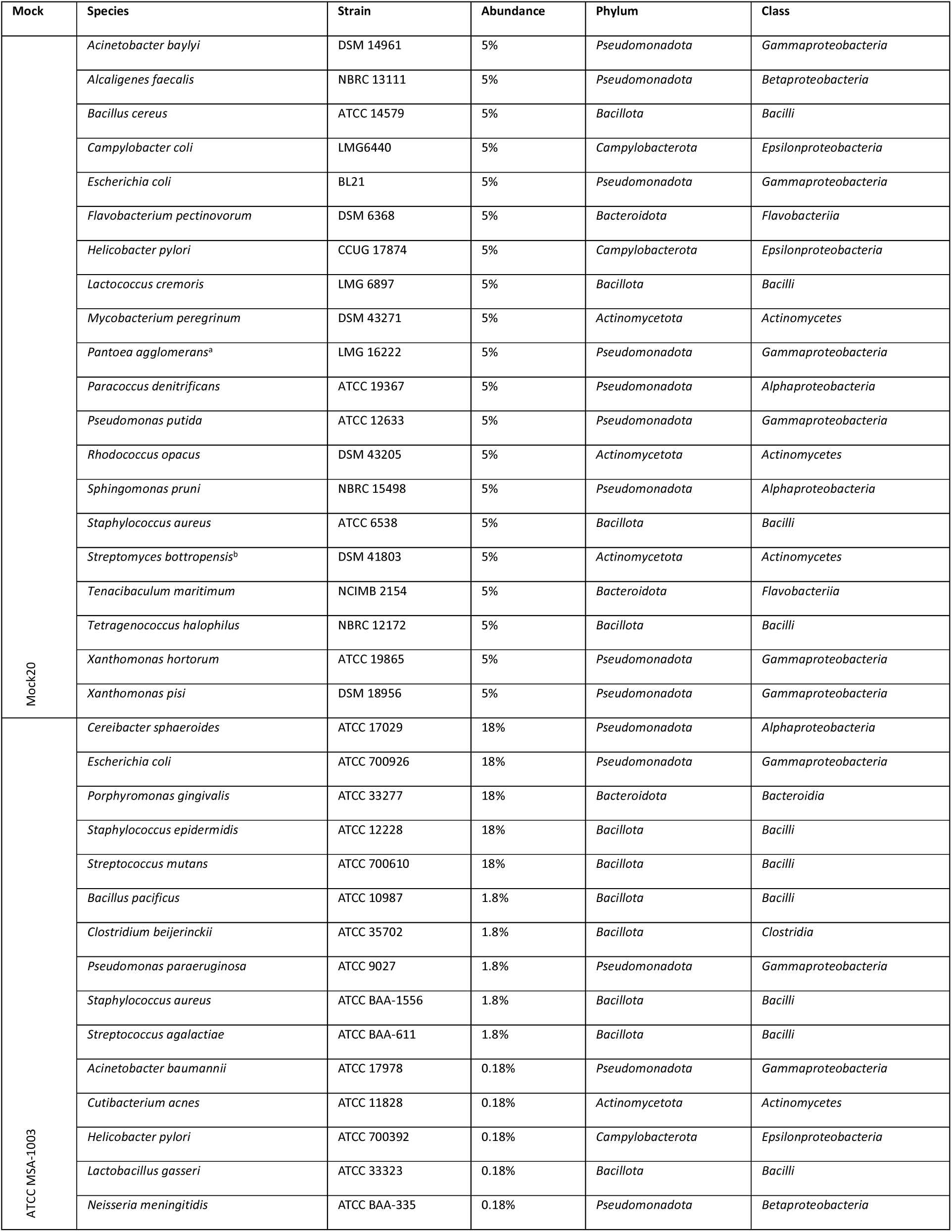
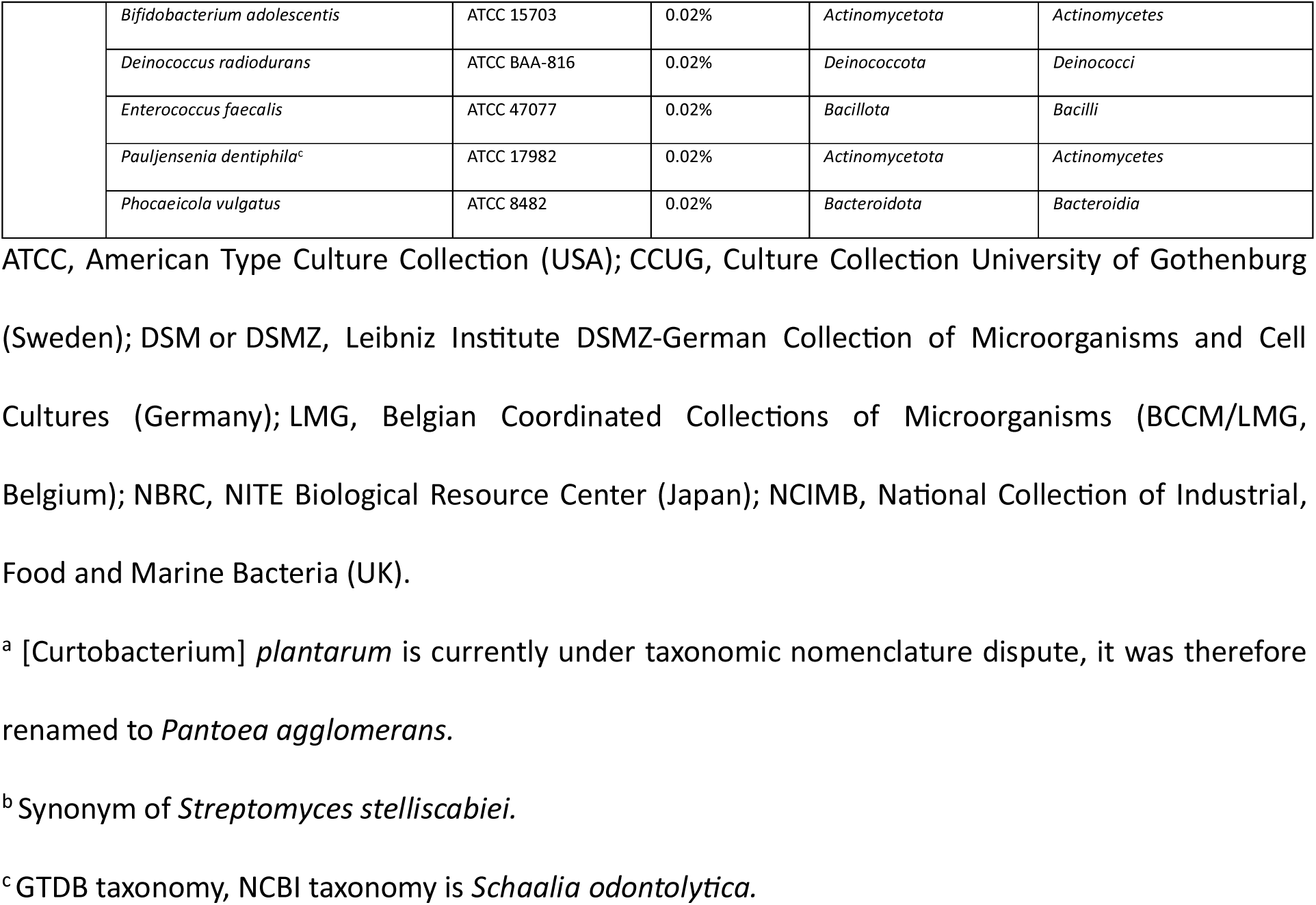
List of bacterial strains within Mock20 and ATCC MSA-1003.

| Mock | Species | Strain | Abundance | Phylum | Class |
| --- | --- | --- | --- | --- | --- |
| Mock20 | *Acinetobacter baylyi* | DSM 14961 | 5% | *Pseudomonadota* | *Gammaproteobacteria* |
|  | *Alcaligenes faecalis* | NBRC 13111 | 5% | *Pseudomonadota* | *Betaproteobacteria* |
|  | *Bacillus cereus* | ATCC 14579 | 5% | *Bacillota* | *Bacilli* |
|  | *Campylobacter coli* | LMG6440 | 5% | *Campylobacterota* | *Epsilonproteobacteria* |
|  | *Escherichia coli* | BL21 | 5% | *Pseudomonadota* | *Gammaproteobacteria* |
|  | *Flavobacterium pectinovorum* | DSM 6368 | 5% | *Bacteroidota* | *Flavobacteriia* |
|  | *Helicobacter pylori* | CCUG 17874 | 5% | *Campylobacterota* | *Epsilonproteobacteria* |
|  | *Lactococcus cremoris* | LMG 6897 | 5% | *Bacillota* | *Bacilli* |
|  | *Mycobacterium peregrinum* | DSM 43271 | 5% | *Actinomycetota* | *Actinomycetes* |
|  | *Pantoea agglomerans*<sup>a</sup> | LMG 16222 | 5% | *Pseudomonadota* | *Gammaproteobacteria* |
|  | *Paracoccus denitrificans* | ATCC 19367 | 5% | *Pseudomonadota* | *Alphaproteobacteria* |
|  | *Pseudomonas putida* | ATCC 12633 | 5% | *Pseudomonadota* | *Gammaproteobacteria* |
|  | *Rhodococcus opacus* | DSM 43205 | 5% | *Actinomycetota* | *Actinomycetes* |
|  | *Sphingomonas pruni* | NBRC 15498 | 5% | *Pseudomonadota* | *Alphaproteobacteria* |
|  | *Staphylococcus aureus* | ATCC 6538 | 5% | *Bacillota* | *Bacilli* |
|  | *Streptomyces bottropensis*<sup>b</sup> | DSM 41803 | 5% | *Actinomycetota* | *Actinomycetes* |
|  | *Tenacibaculum maritimum* | NCIMB 2154 | 5% | *Bacteroidota* | *Flavobacteriia* |
|  | *Tetragenococcus halophilus* | NBRC 12172 | 5% | *Bacillota* | *Bacilli* |
|  | *Xanthomonas hortorum* | ATCC 19865 | 5% | *Pseudomonadota* | *Gammaproteobacteria* |
|  | *Xanthomonas pisi* | DSM 18956 | 5% | *Pseudomonadota* | *Gammaproteobacteria* |
| ATCC MSA-1003 | *Cereibacter sphaeroides* | ATCC 17029 | 18% | *Pseudomonadota* | *Alphaproteobacteria* |
|  | *Escherichia coli* | ATCC 700926 | 18% | *Pseudomonadota* | *Gammaproteobacteria* |
|  | *Porphyromonas gingivalis* | ATCC 33277 | 18% | *Bacteroidota* | *Bacteroidia* |
|  | *Staphylococcus epidermidis* | ATCC 12228 | 18% | *Bacillota* | *Bacilli* |
|  | *Streptococcus mutans* | ATCC 700610 | 18% | *Bacillota* | *Bacilli* |
|  | *Bacillus pacificus* | ATCC 10987 | 1.8% | *Bacillota* | *Bacilli* |
|  | *Clostridium beijerinckii* | ATCC 35702 | 1.8% | *Bacillota* | *Clostridia* |
|  | *Pseudomonas paraeruginosa* | ATCC 9027 | 1.8% | *Pseudomonadota* | *Gammaproteobacteria* |
|  | *Staphylococcus aureus* | ATCC BAA-1556 | 1.8% | *Bacillota* | *Bacilli* |
|  | *Streptococcus agalactiae* | ATCC BAA-611 | 1.8% | *Bacillota* | *Bacilli* |
|  | *Acinetobacter baumannii* | ATCC 17978 | 0.18% | *Pseudomonadota* | *Gammaproteobacteria* |
|  | *Cutibacterium acnes* | ATCC 11828 | 0.18% | *Actinomycetota* | *Actinomycetes* |
|  | *Helicobacter pylori* | ATCC 700392 | 0.18% | *Campylobacterota* | *Epsilonproteobacteria* |
|  | *Lactobacillus gasseri* | ATCC 33323 | 0.18% | *Bacillota* | *Bacilli* |
|  | *Neisseria meningitidis* | ATCC BAA-335 | 0.18% | *Pseudomonadota* | *Betaproteobacteria* |
|  | *Bifidobacterium adolescentis* | ATCC 15703 | 0.02% | *Actinomycetota* | *Actinomycetes* |
|  | *Deinococcus radiodurans* | ATCC BAA-816 | 0.02% | *Deinococcota* | *Deinococci* |
|  | *Enterococcus faecalis* | ATCC 47077 | 0.02% | *Bacillota* | *Bacilli* |
|  | *Pauljensenia dentiphila*<sup>c</sup> | ATCC 17982 | 0.02% | *Actinomycetota* | *Actinomycetes* |
|  | *Phocaeicola vulgatus* | ATCC 8482 | 0.02% | *Bacteroidota* | *Bacteroidia* |

ATCC, American Type Culture Collection (USA); CCUG, Culture Collection University of Gothenburg (Sweden); DSM or DSMZ, Leibniz Institute DSMZ-German Collection of Microorganisms and Cell Cultures (Germany); LMG, Belgian Coordinated Collections of Microorganisms (BCCM/LMG, Belgium); NBRC, NITE Biological Resource Center (Japan); NCIMB, National Collection of Industrial, Food and Marine Bacteria (UK).
<sup>a</sup> [*Curtobacterium*] *plantarum* is currently under taxonomic nomenclature dispute, it was therefore renamed to *Pantoea agglomerans*.
<sup>b</sup> Synonym of *Streptomyces stelliscabiei*.
<sup>c</sup> GTDB taxonomy, NCBI taxonomy is *Schaalia odontolytica*.

Mock20 was constructed by pooling genomic DNA from the 20 strains at equal genome-copy equivalents calculated from genome size and DNA concentration. A staggered commercial mock community, ATCC MSA-1003, consisting of 20 strains ranging from 0.02% to 18% relative abundance, was included as an additional reference standard for method validation and benchmarking (Table 1).

### *rpoB* primer design and *in silico* evaluation

A representative panel of bacterial genomes (N = 17683) was selected from NCBI RefSeq representative species genomes accessed on November 22, 2023 (14); accession numbers are provided in Supplementary File 1. Annotated GenBank files were retrieved for *in silico* gene extraction, primer design, and primer evaluation.

For primer design, *rpoB* sequences were extracted using gene feature annotations (gene=“rpoB”), yielding 14716 sequences. Sequences were aligned with MAFFT v7.520 (15), and conserved regions were identified using Shannon entropy calculated at each alignment position relative to *Escherichia coli* strain K-12 with a 25-position sliding window (16). Universal primer design aimed to maximize cross-taxon coverage while minimizing degeneracy and secondary structure formation.

Primer universality was evaluated by *in silico* PCR using EMBOSS PrimerSearch v6.6.0 (17), allowing up to 20% mismatches per primer. Predicted amplicons were extracted and validated as *rpoB* using HMMER v3.4 (18) with an *rpoB*-specific hidden Markov model constructed from the design alignment. Only HMM-confirmed amplicons were retained as true positives.

### PCR reactions

All primers, sequences, annealing temperatures, extension times, and sources are listed in Table 2. PCR amplifications of rpoB_MV and 16Sfull were performed using Q5 Hot Start High-Fidelity DNA Polymerase (New England Biolabs) in 20 µL reactions containing 0.5 µM of each primer and 5 ng genomic DNA. Cycling consisted of initial denaturation at 98 °C for 1 min; 30 cycles of 98 °C for 10 s, primer-specific annealing for 30 s, and extension at 72 °C; and final extension at 72 °C for 5 min. Amplicon size and quality were verified by agarose gel electrophoresis, and no-template controls were included in all PCRs.

**Table 2.**
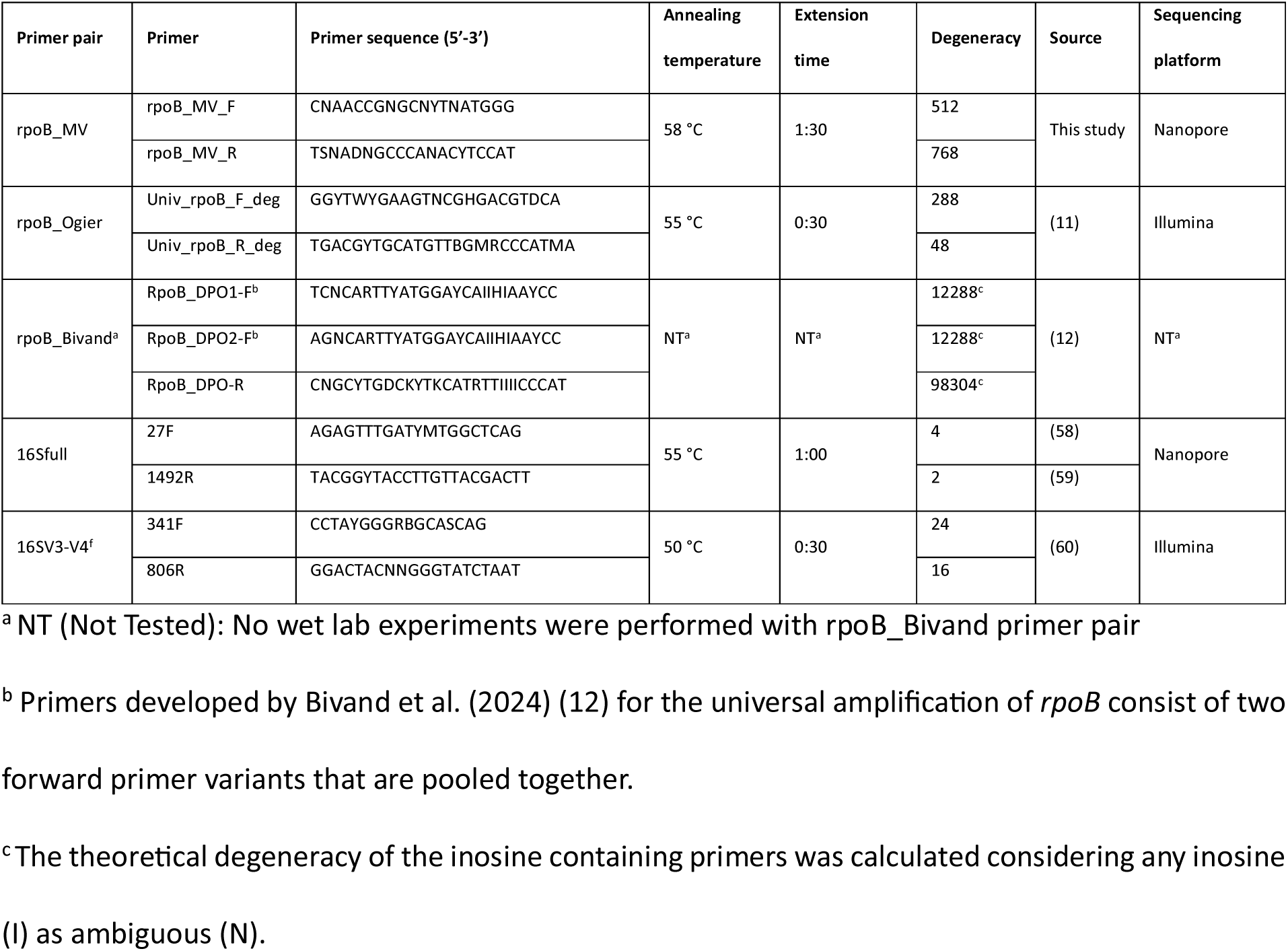
List of primer pairs, corresponding nucleotide sequences, PCR conditions, sources and sequencing platform used.

Amplification of rpoB_Ogier and 16SV3-V4 was performed by Novogene Co., Ltd. (Munich, Germany) using Phusion High-Fidelity PCR Master Mix (Thermo Fisher Scientific), 0.2 µM of each primer, and 10 ng template DNA. Cycling conditions followed the same general program, with primer-specific annealing temperatures listed in Table 2 and 30 s extension at 72 °C.

### Inter-species pairwise distance comparisons

A panel of 287 genera, each containing at least 10 species, was selected from the Genome Taxonomy Database (GTDB) (19). For each species, *rpoB* and 16S rRNA gene sequences were retrieved from GTDB files TIGR02013.fna (20) and bac120_ssu_reps_r226.fna (21). Five primer pairs were evaluated: three targeting *rpoB* and two targeting the 16S rRNA gene (Table 2). Target regions were extracted by *in silico* PCR as described above and aligned per genus and marker using MAFFT v7.520 (15).

Inter-species pairwise distances were calculated for all unique species pairs within each genus and marker. Distances were defined as the number of single nucleotide polymorphisms between aligned sequences, excluding positions where both sequences contained gaps. Mean pairwise distances were calculated per genus, yielding 287 values per marker. Differences between markers were assessed using one-sided Mann-Whitney U tests (alternative hypothesis: greater distances) (22). Marker discrimination was further evaluated using *post hoc* area under the curve (AUC) comparisons, which quantify the probability that a randomly selected inter-species pair exhibits a greater distance for one marker than for another, based on the full distribution of pairwise distances within each genus (23).

### Phylogenetic concordance calculation

Phylogenetic concordance analysis, adapted from (24), was performed for the five genetic markers across the 19 genera represented in Mock20 (Table 1). For each genus, genetic marker-derived phylogenetic trees were compared with corresponding core-genome trees inferred from GTDB representative genomes. Phylogenetic concordance was defined as the percentage of bipartition splits in each marker tree that were congruent with the corresponding core-genome tree.

Core genes were identified with BUSCO v6.6.0 (--auto-lineage) (25), retaining genomes with ≥90% single-copy BUSCO completeness and genes present in ≥95% of genomes within each genus. Core-gene and genetic marker alignments were generated with MAFFT v7.520 (15), trimmed with trimAl v1.5 (26), and used for phylogenetic inference with IQ-TREE v2.3.6 (27) under the GTR model with 1,000 ultrafast bootstrap replicates. Bipartitions with bootstrap support below 50% were collapsed, and phylogenetic concordance was calculated using DendroPy v5.0.8 (28). When multiple genetic marker copies were detected within a genome, one copy was selected at random.

Differences in phylogenetic concordance between markers were assessed using one-sided Mann-Whitney U tests (alternative hypothesis: greater concordance) (22) and *post hoc* AUC comparisons, which quantify the probability that a randomly selected genus exhibits higher concordance for one marker than for another (23).

### Reference database construction

Reference sequence databases were constructed for each primer pair used in the sequencing experiments: 16Sfull, 16SV3-V4, rpoB_MV, and rpoB_Ogier. To enable fair comparison of marker performance, all reference sets were derived from the same GTDB RefSeq genome inputs rather than from marker-specific public databases. Primer-targeted regions were extracted by *in silico* PCR as described above, allowing up to 20% mismatches per primer. Sequences containing ambiguous bases were removed and target identity was confirmed using HMMER v3.4 (18) for *rpoB* genetic markers or barrnap v0.8 (29) for 16S rRNA genetic markers. Validated sequences were clustered at 100% identity using CD-HIT v4.8.1 (30), and GTDB taxonomy was used for taxonomic assignment. Clusters containing identical sequences from multiple taxa were retained as multi-species clusters when species-level discrimination was not possible. Taxonomic naming conflicts at the genus level, likely arising from genomic contamination, were resolved by applying a 90% dominance threshold: if a single genus accounted for more than 90% of sequences within a cluster, it was considered dominant and taxonomies assigned to other genera were treated as contaminants and removed. As GTDB introduces suffixes to distinguish provisional species and genus hypotheses (e.g., Bacillus_A cereus_A and Bacillus_A cereus_B), such labels were normalized to represent the same taxon.

### Sequencing of genetic markers

Each marker was amplified from Mock20 and ATCC MSA-1003 using the corresponding primer pair: rpoB_MV, rpoB_Ogier, 16Sfull, or 16SV3-V4. No-template controls were included for each marker, and amplification products were verified by agarose gel electrophoresis. Amplicons were sequenced using either Oxford Nanopore or Illumina sequencing, depending on marker length and workflow compatibility (Table 2).

### Nanopore sequencing

RpoB_MV and 16Sfull amplicons were purified using AMPure XP magnetic beads (Beckman Coulter), prepared with the SQK-NBD114.24 Native Barcoding Kit (Oxford Nanopore Technologies), and sequenced on an R10.4.1 flow cell using the MinION Mk1B platform (NBA_9168_v114_revP_20Nov2024). Basecalling was performed with Dorado v7.6.7 using the super-accuracy model.

Reads were demultiplexed using Cutadapt v4.7 (31), filtered to retain reads containing both primer sequences, and quality filtered using NanoFilt v2.8.0 (32) with a minimum Q-score of 10. Length filters were set to 1,000-2,300 bp for rpoB_MV and 1,000-1,700 bp for 16Sfull. Filtered reads were aligned to genetic marker-specific reference databases using Minimap2 v2.27 (33) with the map-ont preset. Reads were assigned to the highest-identity reference taxonomy if alignments met ≥90% reference coverage and ≥95% gap-compressed identity.

### Illumina sequencing

Library preparation and sequencing of 16SV3-V4 and rpoB_Ogier amplicons were performed by Novogene Co., Ltd. (Munich, Germany) on an Illumina MiSeq platform using 2 × 300 bp paired-end reads. Demultiplexed reads were processed using VSEARCH v2.27.0 (34). Paired reads were merged, quality filtered, dereplicated, denoised with UNOISE (35), and screened for chimeras using UCHIME3 (36).

ASVs were assigned taxonomy by BLASTn v2.16.0 (37) against the corresponding marker-specific reference database. Assignments were retained when hits met ≥95% sequence identity and ≥90% reference coverage. Equally best hits to different taxa were reported as ambiguous, and ASVs without qualifying hits were left unassigned.

### Accuracy assessment and community profiling

Sequencing results were used to evaluate species-level taxonomic identification and recovery of expected community composition. Relative abundance tables were generated for all downstream analyses. To reduce the impact of cross-contamination, taxa detected in samples where they were not expected were removed unless they represented at least 0.5% of all reads assigned to that taxon within the sequencing run. Filtered counts were then renormalized.

Taxonomic misclassification was quantified as the total relative abundance assigned to species absent from the corresponding mock community (“True Misclassifications”). Reads assigned to species clusters sharing identical reference sequences were classified separately as “Insufficient Resolution”, because reliable species-level discrimination was not possible.

Expected abundance profiles were generated for both mock communities. For 16S rRNA gene-based profiles, expected abundances were corrected for copy number using rrnDB (38), whereas *rpoB* was treated as a single-copy gene. Agreement between observed and expected communities was assessed using Jaccard distance for presence/absence and Bray-Curtis dissimilarity for relative abundance (39, 40).

## RESULTS

### Design of universal primers for long-amplicon *rpoB* amplification

A total of 14716 bacterial *rpoB* sequences from NCBI RefSeq representative genomes were aligned to identify conserved primer-binding sites and variable regions using Shannon entropy analysis (Figure 1). This yielded the rpoB_MV primer pair, which targets conserved regions flanking a variable long-amplicon region.

**Figure 1.**
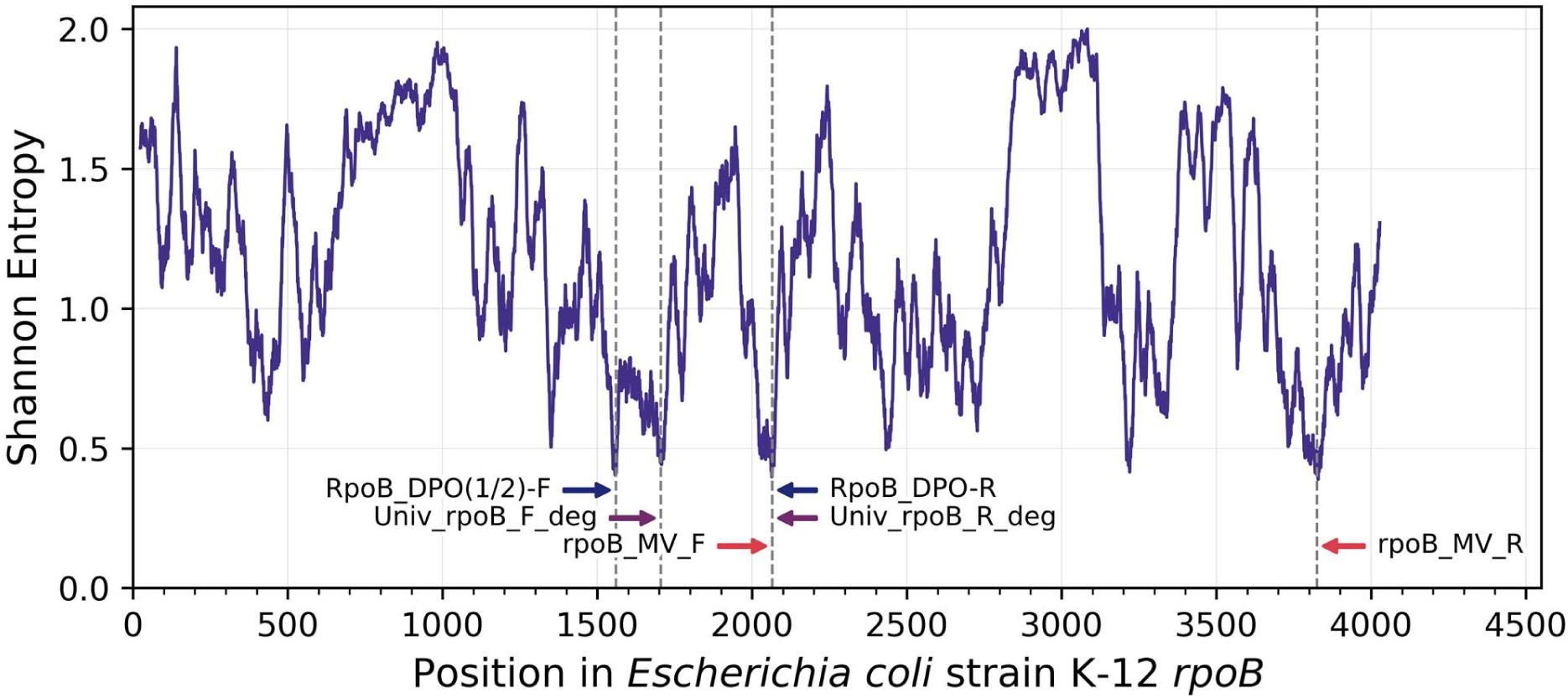
Shannon entropy plot of the *rpoB* genes aligned against *Escherichia coli* strain K-12. Shannon entropy, a measure of sequence variability, was calculated for each nucleotide position in the alignment of *rpoB* genes (N=14716) extracted from the dataset of representative species genomes (N=17683). A moving average window of 25 nucleotides was applied to smooth the entropy profile. Higher Shannon entropy values indicate regions of greater sequence variability, whereas lower values indicate conserved regions. Primer binding sites are indicated by arrows pointing to vertical dashed lines marking the binding positions. Primers designed in this study (rpoB_MV) are indicated in red, whereas primers obtained from literature are shown in purple (rpoB_Bivand) (12), and blue (rpoB_Ogier) (11). Forward and reverse primers for each primer pair are labeled with F and R, respectively.

The corresponding amplicon (rpoB_MV) has a length of 1810 bp in reference to *E. coli* strain K-12. However, a degree of sequence length variability was observed among different bacterial phyla, with per phylum average amplicon sizes ranging from 1356 to 1859 bp (Supplementary File 2; Table S1). Previously reported primer pairs for universal amplification of *rpoB*, rpoB_Ogier and rpoB_Bivand (Table 2), also anneal in conserved regions but only span a region of 434 and 535 bp in reference to *E. coli* strain K-12, respectively.

Preliminary wet-lab PCR validation using the Mock20 panel showed specific amplification of the target fragment in 18 of 20 strains. Amplification failed for *Mycobacterium peregrinum*, whereas *Rhodococcus opacus* produced weak target amplification together with a nonspecific fragment.

### *In silico* evaluation of *rpoB* primer pair universality

Comprehensive *in silico* PCR across 17683 representative bacterial genomes confirmed broad rpoB_MV universality (Figure 2A). A perfect match to both primers was observed in 79.5% of genomes, increasing to 86.2% when one mismatch per primer was allowed. Among the 10 most abundant bacterial phyla in the dataset, reduced predicted universality was observed for *Actinomycetota*, *Mycoplasmatota*, and *Campylobacterota*, with amplification predicted for 62.6%, 34.3%, and 76.9% of genomes, respectively.

**Figure 2.**
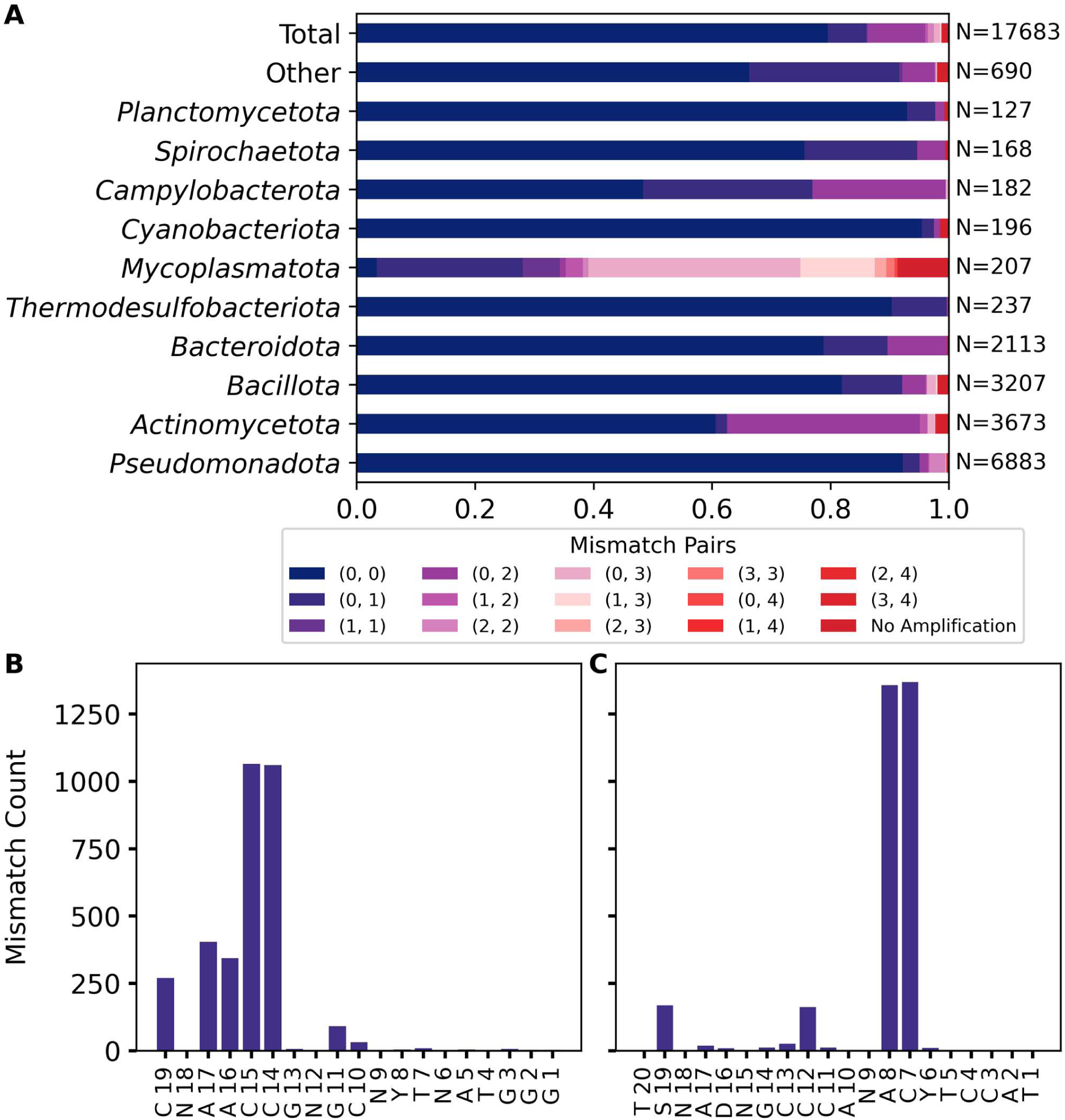
Universality and mismatch analysis of the novel *rpoB* primer pair (rpoB_MV). (A) Bar plot denoting the universality for all bacterial genomes included in this study (N=17683), as well as for the top 10 most abundant phyla. Strains not belonging to the top 10 are categorized in “Other”. In each bar, the possible state of mismatches is presented, in which (X, Y) indicates that one primer has X and the other Y mismatches with the template. The number of bacterial genomes included in each phylum is denoted next to the bar graph. (B-C) Bar plots show the number of mismatches for each nucleotide position (with position 1 being the 3’ end of the primer) in the rpoB_MV forward (B) and reverse primer (C), assessed for all genomes included in the study (N=17683).

Mismatch-position analysis showed that most rpoB_MV_F mismatches occurred near the 5′ end, which are expected to have limited effects on amplification (Figure 2B-C). In contrast, rpoB_MV_R showed elevated mismatch frequencies at the seventh and eighth positions from the 3′ end, primarily in *Actinomycetota*. Applying a conservative criterion that excluded primer pairs with mismatches within four bases of the 3′ end still resulted in 86.2% predicted amplification when one mismatch per primer was allowed.

Compared with published *rpoB* primers, rpoB_MV showed higher predicted universality than rpoB_Ogier and comparable universality to rpoB_Bivand. RpoB_Ogier showed perfect matches in 13.0% of genomes and 34.4% predicted amplification under the above-mentioned conservative mismatch criterion. RpoB_Bivand showed perfect matches in 73.1% of genomes and 93.0% predicted amplification under the same criterion (Supplementary File 3; Figures S1-S2).

### Pairwise distance comparison of the genetic markers

Inter-species pairwise distances were used to compare the discriminatory power of rpoB_MV (i.e., its ability to distinguish between species of the same genus) with 16Sfull, 16SV3-V4, rpoB_Ogier, and rpoB_Bivand genetic markers across 287 genera containing at least 10 species (Figure 3A). Interspecific genetic distances, expressed in the number of SNPs, were calculated for all pairwise combinations of species within a certain genus. Across genera, rpoB_MV showed significantly higher interspecific sequence divergence than both 16Sfull and 16SV3-V4 (p < 0.001), indicating greater species-level discriminatory potential. AUC values for rpoB_MV versus 16Sfull and 16SV3-V4 were 0.94 ± 0.11 and 0.98 ± 0.06, respectively (Supplementary File 3; Figure S3B). RpoB_MV also outperformed the shorter rpoB_Ogier and rpoB_Bivand markers (p < 0.001), with corresponding AUC values of 0.97 ± 0.07 and 0.96 ± 0.08.

**Figure 3.**
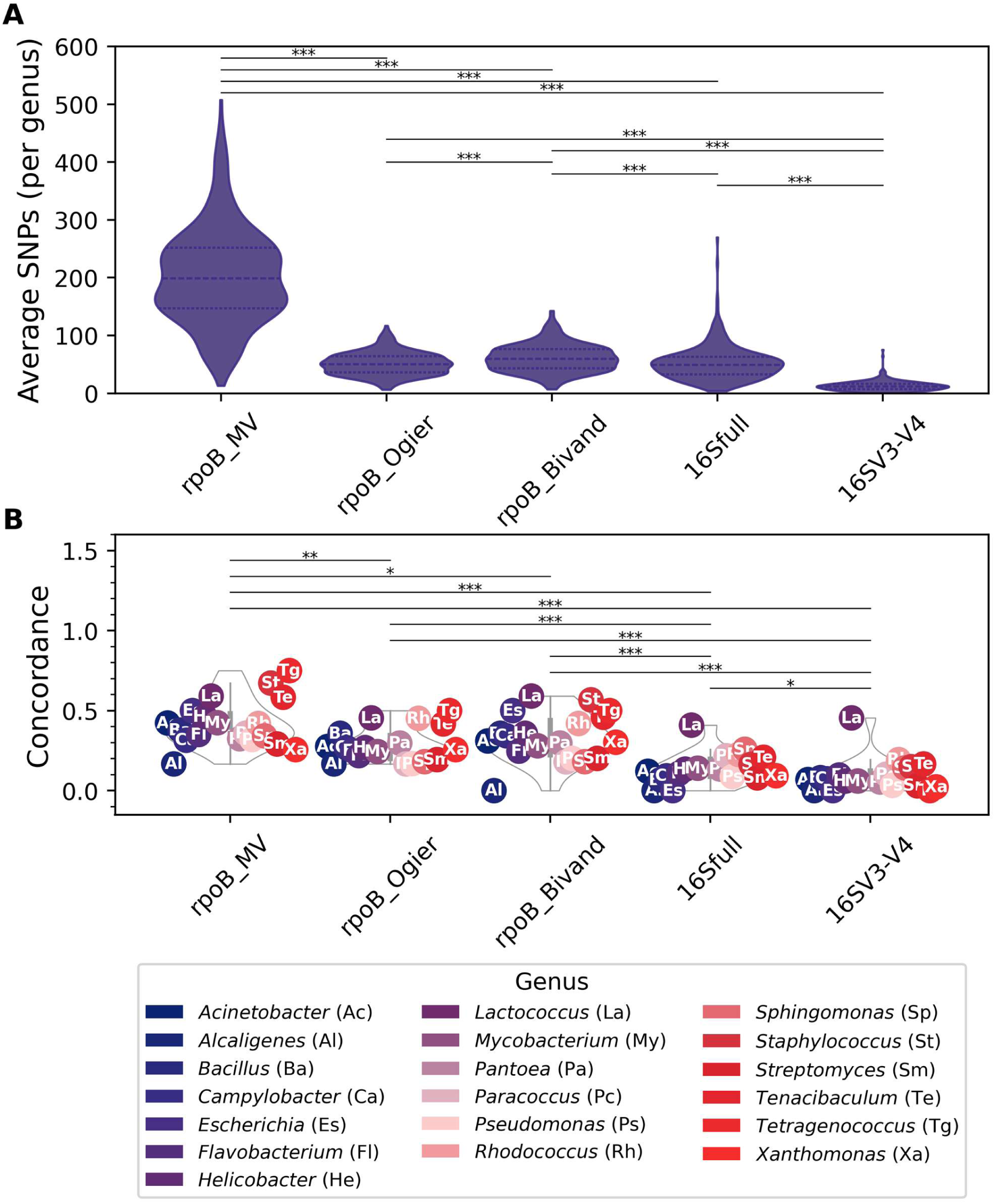
Inter-species distance comparison and phylogenetic concordance analysis of the genetic markers analyzed in this study. (A)Violin plots of inter-species pairwise distances across 287 genera are shown for each of the tested genetic markers. The inter-species pairwise distances (expressed as number of SNPs between two aligned sequences) were calculated for all pair-wise comparisons of unique species within each genus, for which the average number for each genus was used to compile the violin plot. (B) Violin plots show the phylogenetic concordance values calculated for 19 representative genera across the different genetic markers. For each genus, the value is also plotted on top of the violin plot. Significance values from pairwise Mann-Whitney U tests (greater-than) comparing genetic markers based on inter-species distance (A) or phylogenetic concordance (B) across the genera are indicated on the figure (*: *p* <0.05, **: *p* <0.01, ***: *p* <0.001).

### Phylogenetic concordance of the genetic markers

Phylogenetic concordance analysis was performed across the 19 bacterial genera represented in Mock20 to assess whether genetic marker-derived phylogenies reflected core-genome phylogenies (Figure 3B; Supplementary File 2; Table S2). RpoB_MV showed the highest mean concordance among all markers (0.42 ± 0.15), compared with 0.14 ± 0.10 for 16Sfull, 0.09 ± 0.10 for 16SV3-V4, 0.33 ± 0.15 for rpoB_Bivand, and 0.29 ± 0.11 for rpoB_Ogier. Concordance could not be calculated for 16Sfull in *Tetragenococcus* because full-length 16S rRNA gene sequences were absent from several genomes, or for rpoB_Ogier in *Staphylococcus* because of excessive forward-primer mismatches.

Overall, rpoB_MV significantly outperformed both 16S rRNA gene markers (p < 0.001; AUC = 0.96 for both comparisons) and the previously reported *rpoB* markers (p < 0.05; AUC = 0.78 versus rpoB_Ogier and 0.66 versus rpoB_Bivand) (Supplementary File 3; Figure S4B).

### Assessing the performance of *rpoB*-based primers for metataxonomic profiling via amplicon sequencing of mock communities

The performance of rpoB_MV was evaluated using nanopore sequencing of two mock communities: the in-house equimolar Mock20 community and the staggered ATCC MSA-1003 community. Performance was benchmarked against full-length 16S rRNA gene nanopore sequencing, 16S V3-V4 Illumina sequencing, and rpoB_Ogier Illumina sequencing. Two technical replicates were analyzed for each genetic marker and mock community. Sequencing results are summarized in Supplementary File 2; Table S3, and relative abundance tables are provided in Supplementary File 2; Tables S4-S7.

### Evaluation of the primer pairs for species-level classifications

RpoB_MV detected all species in Mock20 and 18 of 20 species in ATCC MSA-1003, missing *Clostridium beijerinckii* and *Pauljensenia dentiphila* (Figure 4). Misclassification rates were extremely low, with 0.007 ± 0.000% of reads incorrectly assigned in Mock20 and 0.001 ± 0.000% in ATCC MSA-1003 (True Misclassifications). No rpoB_MV reads were assigned to unresolved species clusters (Insufficient Resolution).

**Figure 4.**
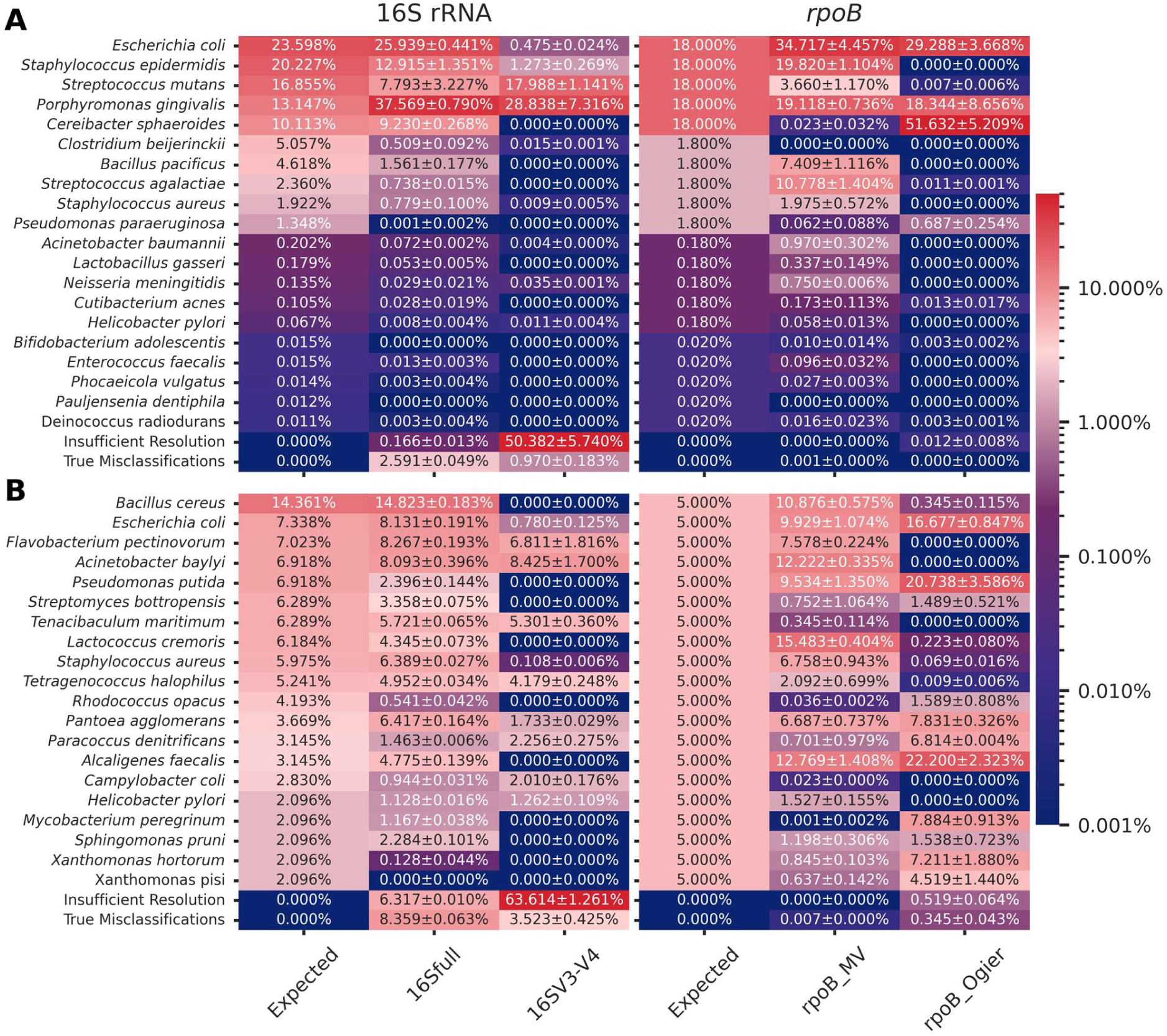
Heat map of the relative abundance of taxa detected using the different primer pairs across two mock communities. The average relative abundance of two technical replicates +/- the standard deviation is shown for the ATCC MSA-1003 (A) and the Mock20 (B) mock community. The category “True Misclassifications” reflects the percentage of reads assigned to species not present in the mock community, while category “Insufficient Resolution” reflects the percentage of reads assigned to species clusters (i.e., impossible to differentiate based on the genetic marker used). The color scale is logarithmic and represents the relative abundance of the taxa identified, ranging from 0.001% to 50%.

RpoB_Ogier failed to identify 5 taxa in Mock20 and 11 taxa in ATCC MSA-1003. Misclassification rates remained low, at 0.345 ± 0.043% for Mock20 and 0.000 ± 0.000% for ATCC MSA-1003, while unresolved species-cluster assignments accounted for 0.519 ± 0.064% and 0.012 ± 0.008% of reads, respectively.

The 16S rRNA genetic markers showed lower species-level performance. Full-length 16S failed to detect *Xanthomonas pisi* in Mock20 and two low-abundance species in ATCC MSA-1003 (*Bifidobacterium adolescentis* and *P. dentiphila*), with misclassification rates of 8.359 ± 0.063% and 2.591 ± 0.049%, respectively, while 0.166 ± 0.013% and 6.317 ± 0.010% of the obtained reads were assigned to unresolved species clusters. The 16SV3-V4 marker detected only 10 species in Mock20 and 9 species in ATCC MSA-1003, and a large fraction of reads were assigned to unresolved species clusters: 63.614 ± 1.261% and 50.382 ± 5.740%, respectively. Misclassification rates were 3.523 ± 0.425% for Mock20 and 0.970 ± 0.183% for ATCC MSA-1003.

### Evaluation of community composition accuracy

Community composition accuracy was evaluated using log_10_-fold deviation, Jaccard distance, and Bray-Curtis dissimilarity (Figure 5; Supplementary File 3; Figure S5). Using a log_10_-fold deviation threshold of ≤−1, rpoB_MV severely underrepresented 9 taxa across both mock communities, compared with 23 taxa for rpoB_Ogier, 6 for 16Sfull, and 27 for 16SV3-V4.

**Figure 5.**
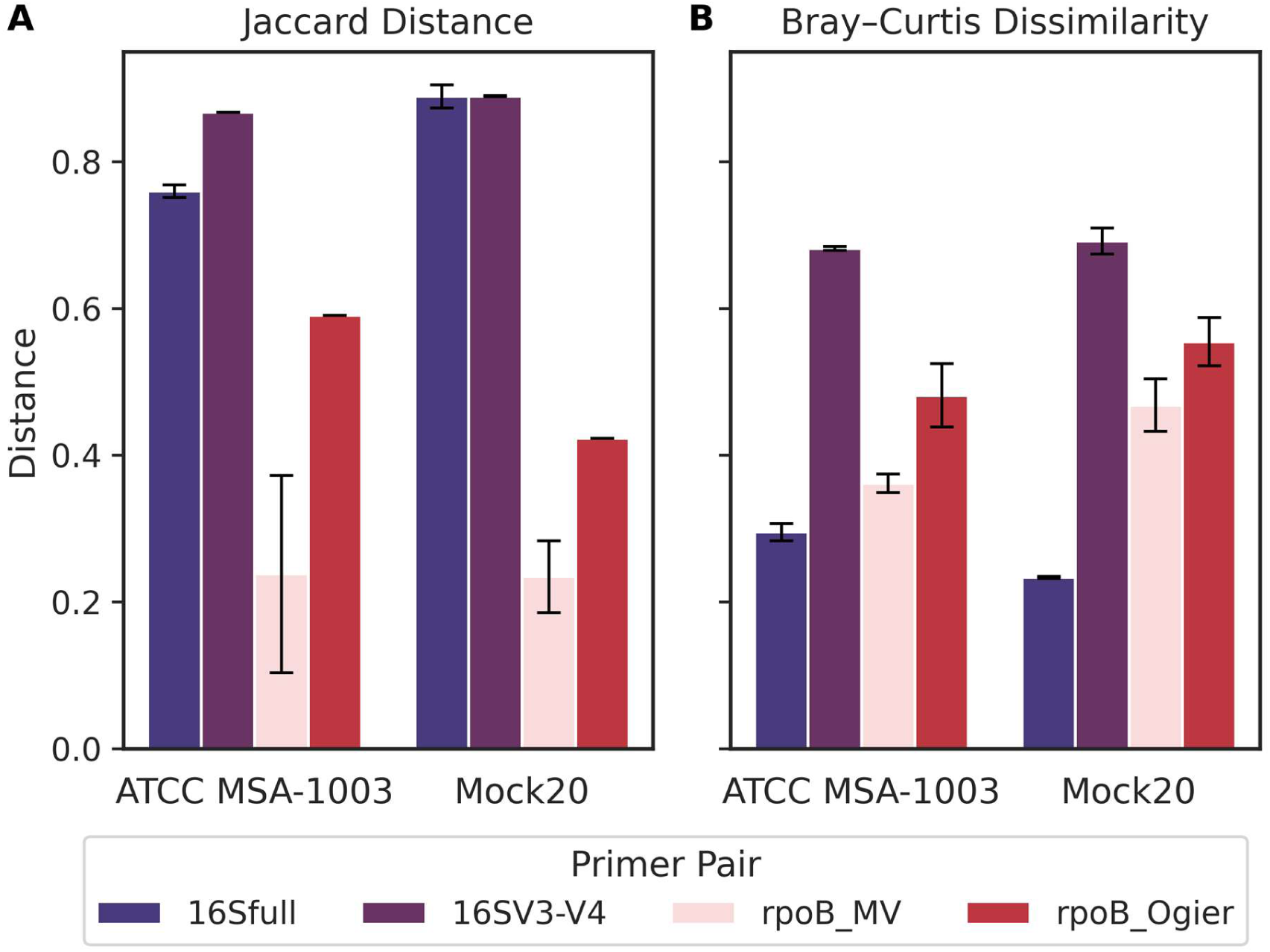
Jaccard distance and Bray-Curtis dissimilarity for the different genetic markers across two mock communities. Both the Jaccard distance (A) and the Bray-Curtis dissimilarity (B) range from 0 to 1, with values closer to 0 indicating greater similarity between observed and expected community compositions, and a value of 1 representing complete divergence. The average values and their standard deviations are shown for the two technical replicates performed.

RpoB_MV produced the lowest Jaccard distances among all markers, with values of 0.23 ± 0.05 for Mock20 and 0.24 ± 0.13 for ATCC MSA-1003 (Figure 5A). In comparison, Jaccard distances were higher for rpoB_Ogier, 16Sfull, and 16SV3-V4, indicating lower presence/absence agreement with expected species composition.

When relative abundance was considered, rpoB_MV showed lower Bray-Curtis dissimilarity than rpoB_Ogier and 16SV3-V4, with values of 0.47 ± 0.04 for Mock20 and 0.36 ± 0.01 for ATCC MSA-1003 (Figure 5B). However, 16Sfull yielded the lowest Bray-Curtis dissimilarities overall, suggesting higher quantitative accuracy despite poorer species-level detection.

## DISCUSSION

The rpoB_MV primers were designed based on conserved regions identified by low Shannon entropy values (Figure 1). Effective universal primer design requires minimizing primer-template mismatches, particularly near the 3′ end, where mismatches severely reduce PCR efficiency (41–44). For protein-encoding genes, degenerate primers are necessary to accommodate sequence variability and achieve broad bacterial coverage. However, increasing degeneracy reduces the effective concentration of the most suited primers and can compromise amplification efficiency and specificity, particularly when degenerate positions occur near the 3′ end (45, 46). The rpoB_MV primers balance these constraints with total degeneracies of 512 (forward) and 768 (reverse), while degenerate positions and mismatches near the 3′ end were avoided. Evaluation across 20 bacterial species, spanning multiple phyla, yielded specific amplification for 18 strains (90%). While *R. opacus* and *M. peregrinum*, each carrying two reverse-primer mismatches, failed amplification, *Campylobacter coli* amplified successfully despite a single forward-primer mismatch. A conservative universality threshold was therefore defined: at most one mismatch per primer, with none within four nucleotides of the 3′ end. A comprehensive *in silico* PCR across 17683 reference genomes confirmed broad universality (Figure 2): over 86% of the tested strains were predicted to amplify under the experimentally derived thresholds. Three phyla showed reduced universality caused by increased mismatch levels. In *Actinomycetota*, to which *R. opacus* and *M. peregrinum* belong, a double mismatch in rpoB_MV_R (positions 7-8 from the 3′ end) affected ∼27% of genomes (N = 3673). *Campylobacterota* (N = 182) and *Mycoplasmatota* (N = 207) also showed reduced universality but each represents only ∼1% of the dataset. Compared with published *rpoB* primers under identical criteria, rpoB_Ogier showed predicted amplification for only 34.5% of genomes, while rpoB_Bivand reached 93.0% (Supplementary File 3; Figure S1). However, rpoB_Bivand generates a much shorter amplicon (∼535 bp vs. ∼1,810 bp) and relies on inosine bases, which form less stable duplexes with variable binding strengths (47), are subject to nearest-neighbor effects (48), and markedly increase primer degeneracy (Table 2). This likely introduces amplification bias and may distort community composition estimates. Additionally, inosine-containing primers are poorly compatible with proofreading polymerases (49, 50).

Beyond broad amplification, the performance of a genetic marker also depends on its ability to differentiate taxa. Species-level identification is essential for accurately inferring ecological functions and pathogenic potential of microbial community members, yet the 16S rRNA gene often lacks sufficient discriminatory power to distinguish closely related species, such as within *Bacillus* (8). Protein-encoding housekeeping genes like *rpoB* offer greater sequence variability and more reliable species-level resolution (8, 9). Two complementary metrics were used to evaluate the discriminative power of the rpoB_MV amplicon. First, inter-species pairwise nucleotide distances across 287 genera showed that rpoB_MV exhibits substantially higher interspecific sequence divergence than both 16Sfull and 16SV3-V4 (Figure 3A), which is in agreement with the higher evolutionary rate of *rpoB* (8, 9, 24). The larger amplicon size further increases the absolute number of informative positions compared with shorter *rpoB* markers, even when relative divergence per unit length is comparable. Second, phylogenetic concordance with core-genome trees was assessed across the 19 genera in Mock20 (Figure 3B). RpoB_MV showed significantly higher concordance than both 16S rRNA genetic markers and the shorter *rpoB* genetic markers, indicating that its sequence variation more reliably reflects evolutionary relationships. This is consistent with Hassler *et al.* (2022) (24), who showed that 16S rRNA phylogenies often diverge from core-genome trees due to high conservation and susceptibility to horizontal gene transfer. Furthermore, they demonstrated that phylogenetic concordance increases with the number of informative positions, which likely explains the improved performance of rpoB_MV over the shorter *rpoB* markers. Using a genetic marker with a strong phylogenetic signal reduces the risk of systematic misclassification (51). Altogether, these findings indicate that the rpoB_MV genetic marker combines high sequence divergence with strong evolutionary coherence, making it a suitable tool for accurate species-level identification in metataxonomic studies.

In addition to the extensive *in silico* analyses, species-level accuracy was evaluated by benchmarking the rpoB_MV sequencing workflow against 16Sfull, 16SV3-V4, and rpoB_Ogier using equimolar (Mock20) and staggered (ATCC MSA-1003) mock communities, assessing taxon recovery, correctness of species assignments, and false positive/negative rates. RpoB_MV recovered all expected species in Mock20 and 18/20 in ATCC MSA-1003 (Figure 4). The two undetected species were *C. beijerinckii* (four forward-primer mismatches) and *P. dentiphila*, a low-abundance and GC-rich taxon (0.02% abundance, 67.5% GC). Notably, *R. opacus* and *M. peregrinum* were detected at low abundance in Mock20 despite producing no visible amplicons during PCR evaluation. Similarly, *B. adolescentis* and *Cutibacterium acnes*, each with two forward-primer mismatches near the 5′ end, were successfully detected in ATCC MSA-1003. True misclassification rates (reads assigned to species absent from the mock community) were exceptionally low for rpoB_MV (≤0.007%), with the few misclassified reads corresponding to taxa from the other mock community, suggesting low-level cross-contamination or barcode cross-talk (52). No rpoB_MV reads were assigned to unresolved species clusters. RpoB_Ogier also showed limited misclassification (≤0.35%) but failed to detect multiple species, likely due to primer-template mismatches (Supplementary File 3; Figures S1B and S2B), and a small fraction of reads were assigned to species clusters. Both 16S markers exhibited substantially higher misclassification rates (≤8.4% for 16Sfull; ≤3.5% for 16SV3-V4), and a large fraction of 16SV3-V4 reads, and to a lesser extent 16Sfull reads, were assigned to unresolved species clusters (Figure 4; Supplementary File 2; Tables S4 and S6), reflecting the limited taxonomic resolution of these markers. The higher misclassification rates observed for 16Sfull may be partly exacerbated by nanopore sequencing errors, which are more likely to result in incorrect species assignments when inter-species sequence divergence is low. Similarly, rpoB_MV yielded the lowest Jaccard distances (Figure 5A), reflecting high species recovery with minimal false positives, whereas both 16S rRNA genetic markers showed markedly higher Jaccard distances due to misclassifications and incomplete recovery. Although full-length 16S rRNA sequencing improves resolution over V3-V4 (6, 7), it still failed to distinguish closely related species. For example, *X. pisi* could not be resolved from other *Xanthomonas* species using either 16S rRNA genetic marker, whereas rpoB_MV enabled reliable identification (Supplementary File 2; Table S6). These results align with the pairwise distance analyses. For example, rpoB_MV yielded 143 average SNPs versus 20 for 16Sfull within *Xanthomonas*. The extremely low misclassification rates observed for rpoB_MV contrast with short-read 16S rRNA gene sequencing approaches, where abundance thresholds are often applied to mitigate false positive identifications, enabling more confident detection of low-abundance taxa.

Although rpoB_MV demonstrated high species-level identification accuracy, it was prone to relative abundance bias. This was assessed using log₁₀-fold deviation between observed and expected relative abundance, and Bray-Curtis dissimilarity. Across both mock communities, 9 taxa were underrepresented more than 10-fold with rpoB_MV, compared with 5 for 16Sfull, 27 for 16SV3-V4, and 23 for rpoB_Ogier (Supplementary File 3; Figure S5). The high number for 16SV3-V4 should be interpreted cautiously, as it likely reflects reads assigned to species clusters rather than amplification failure. For overall community composition accuracy, rpoB_MV was slightly outperformed by 16Sfull in Bray-Curtis dissimilarity (Figure 5), particularly for Mock20. This likely reflects the greater sensitivity of equimolar communities to missing taxa: *R. opacus* and *M. peregrinum* were strongly underrepresented in Mock20, substantially influencing dissimilarity, whereas the undetected low-abundance taxa in ATCC MSA-1003 (*C. beijerinckii* and *P. dentiphila*) had less impact. RpoB_Ogier ranked third, and 16SV3-V4 performed worst. It should be noted that expected abundances for the 16S rRNA genetic markers were corrected for gene copy number using rrnDB (38), whereas *rpoB*, as a single-copy gene, required no such correction. While this correction improves quantitative accuracy for 16S-based markers, it relies on the availability and accuracy of copy number estimates, which may not be complete for all taxa. Underrepresentation in rpoB_MV samples could be attributed to two main factors. First, underrepresentation could be caused by primer-template mismatches, as observed for *C. beijerinckii*, which carries 4 forward-primer mismatches, while *M. peregrinum* and *R. opacus* each have two reverse-primer mismatches. Second, extreme GC-content (<35% or >65%) was associated with severe underrepresentation, as observed for *S. bottropensis* (67.7%) or *T. maritimum* (34.7%), as well as increased inter-replicate variability, consistent with previous reports (53, 54). Although amplicon length variation between taxa cannot be excluded as a contributing factor (55), multiple regression analysis confirmed GC-content and mismatch count as the strongest drivers of amplification bias, with no significant effect of amplicon length (Supplementary File 2; Table S8).

Despite the clear improvement of species-level classifications using rpoB_MV, some limitations must be acknowledged. Although high universality was achieved, rpoB_MV is unlikely to match the universality of 16S rRNA gene primers due to silent third-codon mutations inherent to protein-encoding genes (10). Universality was especially reduced in *Actinomycetota*, due to a double mismatch in the reverse primer of ∼27% of the genomes. Increasing primer degeneracy to accommodate these mismatches would reach impractical levels. Instead, we suggest pooling the current primers with a supplementary reverse-primer (5’-TGYAWBGCCCARCAYTCCAT-3’, degeneracy 48) with high complementarity to *Actinomycetota*, though PCR optimization would be required. GC-content bias, a common challenge in amplicon sequencing (54), also affected rpoB_MV performance. This could potentially be mitigated by increasing initial denaturation times or slowing annealing ramp rates (53). Inter-replicate variability due to extreme GC% may be reduced by pooling PCR replicates before sequencing (56, 57). Given moderate amplification bias, a complementary strategy combining rpoB_MV with 16S rRNA gene sequencing could balance species-level resolution with quantitative robustness.

In conclusion, this study presents the rpoB_MV primer pair, a highly universal primer set targeting an approximately 1.8 kb region of the bacterial *rpoB* gene for long-read metataxonomic profiling. The rpoB_MV amplicon enabled substantially higher species-level identification accuracy than conventional 16S rRNA genetic markers, with comparable species recovery. These characteristics make the method particularly valuable in settings where species-level accuracy directly affects biological interpretation, such as tracking foodborne pathogens in complex microbial communities, resolving functionally distinct species in plant-associated microbiomes, or identifying clinically relevant organisms in diagnostic workflows. The close agreement between sequencing outcomes and *in silico* divergence analyses further confirms that computational approaches are effective tools for assessing the discriminative potential of genetic markers. The integrated primer design and evaluation framework provides a generalizable approach for developing universal primers. To our knowledge, this is the first broadly validated universal primer set for long-read *rpoB* metataxonomics, representing a major advance for high-resolution microbiome research and diagnostic applications.

## Acknowledgements

Part of the computational resources and services used in this work were provided by the VSC (Flemish Supercomputer Center), funded by the Research Foundation - Flanders (FWO) and the Flemish Government.

## Data, Metadata, and Code Availability

The sequencing data generated in this study have been deposited in the European Nucleotide Archive (ENA) at EMBL-EBI under the study accession PRJEB109242 and are publicly available at: https://www.ebi.ac.uk/ena/browser/view/PRJEB109242.

The reference databases generated in this study and the scripts used in this study have been deposited on Zenodo and are publicly available at: 10.5281/zenodo.18941399.

## Authors’ contributions

MV, HR, SC conceived the concept and structure of the study. The initial draft was written by MV and HR. MV performed the primer design, *in silico* performance assessments, and wet-lab experiments. SC and MV performed the Illumina and nanopore sequencing analysis, respectively. All authors contributed to writing. Final revision was done by MV, HR, SC. All authors read and approved the final manuscript. HR provided funding.

## Supplementary Files

Supplementary File 1

Text file (.txt)

NCBI genome accession identifiers used for primer design and universality assessment.

Supplementary File 2

Excel file (.xls)

Table S1. Amplicon lengths for the given primer pairs across the 10 most abundant phyla in the genome dataset.

Table S2. Phylogenetic concordance results for each genus and primer pair used in this study.

Table S3. Summary of the sequencing output for the different samples.

Table S4. Relative abundance table for ATCC MSA-1003, with primers targeting the 16S rRNA gene.

Table S5. Relative abundance table for ATCC MSA-1003, with primers targeting *rpoB*.

Table S6. Relative abundance table for Mock20, with primers targeting the 16S rRNA gene.

Table S7. Relative abundance table for Mock20, with primers targeting *rpoB*.

Table S8. Multiple regression analysis of the observed bias (log₂-transformed) in terms of GC-content, amplicon length, and total mismatches.

Supplementary File 3

PDF file (.pdf)

Figure S1. Universality analysis of rpoB primers utilized in this study. Figure S2. Mismatch position profiles of the rpoB primer pairs (rpoB_MV, rpoB_Ogier, and rpoB_Bivand) against the genome dataset.

Figure S3. Inter-species distance comparison of the primer pairs analyzed in this study.

Figure S4. Phylogenetic concordance analysis of the primer pairs analyzed in this study.

Figure S5. Heatmap of the log₁₀-fold deviation between observed and expected taxon abundances in the mock communities.

## References

1. Venbrux M, Crauwels S, Rediers H. 2023. Current and emerging trends in techniques for plant pathogen detection. Front Plant Sci 14.

2. Johnson JS, Spakowicz DJ, Hong B-Y, Petersen LM, Demkowicz P, Chen L, Leopold SR, Hanson BM, Agresta HO, Gerstein M, Sodergren E, Weinstock GM. 2019. Evaluation of 16S rRNA gene sequencing for species and strain-level microbiome analysis. Nat Commun 10.

3. Petrone JR, Rios Glusberger P, George CD, Milletich PL, Ahrens AP, Roesch LFW, Triplett EW. 2023. RESCUE: a validated Nanopore pipeline to classify bacteria through long-read, 16S-ITS-23S rRNA sequencing. Front Microbiol 14.

4. Abellan-Schneyder I, Matchado MS, Reitmeier S, Sommer A, Sewald Z, Baumbach J, List M, Neuhaus K. 2021. Primer, Pipelines, Parameters: Issues in 16S rRNA Gene Sequencing. mSphere 6:10.1128/msphere.01202-20.

5. Vasileiadis S, Puglisi E, Arena M, Cappa F, Cocconcelli PS, Trevisan M. 2012. Soil Bacterial Diversity Screening Using Single 16S rRNA Gene V Regions Coupled with Multi-Million Read Generating Sequencing Technologies. PLoS ONE 7:e42671.

6. Esberg A, Fries N, Haworth S, Johansson I. 2024. Saliva microbiome profiling by full-gene 16S rRNA Oxford Nanopore Technology versus Illumina MiSeq sequencing. npj Biofilms Microbiomes 10:149, s41522-024-00634–1.

7. Matsuo Y, Komiya S, Yasumizu Y, Yasuoka Y, Mizushima K, Takagi T, Kryukov K, Fukuda A, Morimoto Y, Naito Y, Okada H, Bono H, Nakagawa S, Hirota K. 2021. Full-length 16S rRNA gene amplicon analysis of human gut microbiota using MinION^TM^ nanopore sequencing confers species-level resolution. BMC Microbiology 21:35.

8. Reller LB, Weinstein MP, Petti CA. 2007. Detection and Identification of Microorganisms by Gene Amplification and Sequencing. 8. Clinical Infectious Diseases 44:1108–1114.

9. Adékambi T, Drancourt M, Raoult D. 2009. The rpoB gene as a tool for clinical microbiologists. 1. Trends in Microbiology 17:37–45.

10. Case RJ, Boucher Y, Dahllöf I, Holmström C, Doolittle WF, Kjelleberg S. 2007. Use of 16S rRNA and *rpoB* Genes as Molecular Markers for Microbial Ecology Studies. Appl Environ Microbiol 73:278–288.

11. Ogier J-C, Pagès S, Galan M, Barret M, Gaudriault S. 2019. rpoB, a promising marker for analyzing the diversity of bacterial communities by amplicon sequencing. BMC Microbiol 19:171.

12. Bivand JM, Dyrhovden R, Sivertsen A, Tellevik MG, Patel R, Kommedal Ø. 2024. Broad-range amplification and sequencing of the rpoB gene: a novel assay for bacterial identification in clinical microbiology. J Clin Microbiol 62:e00266–24.

13. Vos M, Quince C, Pijl AS, Hollander M de, Kowalchuk GA. 2012. A Comparison of rpoB and 16S rRNA as Markers in Pyrosequencing Studies of Bacterial Diversity. PLOS ONE 7:e30600.

14. Goldfarb T, Kodali VK, Pujar S, Brover V, Robbertse B, Farrell CM, Oh D-H, Astashyn A, Ermolaeva O, Haddad D, Hlavina W, Hoffman J, Jackson JD, Joardar VS, Kristensen D, Masterson P, McGarvey KM, McVeigh R, Mozes E, Murphy MR, Schafer SS, Souvorov A, Spurrier B, Strope PK, Sun H, Vatsan AR, Wallin C, Webb D, Brister JR, Hatcher E, Kimchi A, Klimke W, Marchler-Bauer A, Pruitt KD, Thibaud-Nissen F, Murphy TD. 2025. NCBI RefSeq: reference sequence standards through 25 years of curation and annotation. Nucleic Acids Research 53:D243–D257.

15. Kuraku S, Zmasek CM, Nishimura O, Katoh K. 2013. aLeaves facilitates on-demand exploration of metazoan gene family trees on MAFFT sequence alignment server with enhanced interactivity. Nucleic Acids Research 41:W22–W28.

16. Shannon CE. 1948. A mathematical theory of communication. The Bell System Technical Journal 27:379–423.

17. Rice P, Longden I, Bleasby A. 2000. EMBOSS: the European Molecular Biology Open Software Suite. 6. Trends Genet 16:276–277.

18. Eddy SR. 2011. Accelerated Profile HMM Searches. PLoS Comput Biol 7:e1002195.

19. Parks DH, Chuvochina M, Rinke C, Mussig AJ, Chaumeil P-A, Hugenholtz P. 2022. GTDB: an ongoing census of bacterial and archaeal diversity through a phylogenetically consistent, rank normalized and complete genome-based taxonomy. Nucleic Acids Research 50:D785–D794.

20. 2025. Genome Taxonomy Database (GTDB). bac120_marker_genes_reps_r226.tar.gz (Release 226.0).

21. 2025. Genome Taxonomy Database (GTDB). bac120_ssu_reps_r226.fna.gz (Release 226.0).

22. Mann HB, Whitney DR. 1947. On a Test of Whether one of Two Random Variables is Stochastically Larger than the Other. Ann Math Statist 18:50–60.

23. Hanley JA, McNeil BJ. 1983. A method of comparing the areas under receiver operating characteristic curves derived from the same cases. Radiology 148:839–843.

24. Hassler HB, Probert B, Moore C, Lawson E, Jackson RW, Russell BT, Richards VP. 2022. Phylogenies of the 16S rRNA gene and its hypervariable regions lack concordance with core genome phylogenies. Microbiome 10:104.

25. Tegenfeldt F, Kuznetsov D, Manni M, Berkeley M, Zdobnov EM, Kriventseva EV. 2025. OrthoDB and BUSCO update: annotation of orthologs with wider sampling of genomes. Nucleic Acids Research 53:D516–D522.

26. Capella-Gutiérrez S, Silla-Martinez JM, Gabaldón T. 2009. trimAl: a tool for automated alignment trimming in large-scale phylogenetic analyses. Bioinformatics 25:1972–1973.

27. Nguyen L-T, Schmidt HA, Von Haeseler A, Minh BQ. 2015. IQ-TREE: A Fast and Effective Stochastic Algorithm for Estimating Maximum-Likelihood Phylogenies. Molecular Biology and Evolution 32:268–274.

28. Moreno MA, Holder MT, Sukumaran J. 2024. DendroPy 5: a mature Python library for phylogeneticcomputing. JOSS 9:6943.

29. Seemann T. Barrnap 0.8: rapid ribosomal RNA prediction.

30. Li W, Godzik A. 2006. Cd-hit: a fast program for clustering and comparing large sets of protein or nucleotide sequences. Bioinformatics 22:1658–1659.

31. Martin M. 2011. Cutadapt removes adapter sequences from high-throughput sequencing reads. EMBnet j 17:10.

32. Coster WD. 2017. Nanofilt. Python.

33. Li H. 2021. New strategies to improve minimap2 alignment accuracy. Bioinformatics 37:4572–4574.

34. Rognes T, Flouri T, Nichols B, Quince C, Mahé F. 2016. VSEARCH: a versatile open source tool for metagenomics. PeerJ 4:e2584.

35. Edgar RC. 2016. UNOISE2: improved error-correction for Illumina 16S and ITS amplicon sequencing 10.1101/081257.

36. Edgar RC, Haas BJ, Clemente JC, Quince C, Knight R. 2011. UCHIME improves sensitivity and speed of chimera detection. Bioinformatics 27:2194–2200.

37. Camacho C, Coulouris G, Avagyan V, Ma N, Papadopoulos J, Bealer K, Madden TL. 2009. BLAST+: architecture and applications. BMC Bioinformatics 10:421.

38. Klappenbach JA. 2001. rrndb: the Ribosomal RNA Operon Copy Number Database. Nucleic Acids Research 29:181–184.

39. Jaccard P. 1912. THE DISTRIBUTION OF THE FLORA IN THE ALPINE ZONE.^1^. New Phytologist 11:37–50.

40. Bray JR, Curtis JT. 1957. An Ordination of the Upland Forest Communities of Southern Wisconsin. Ecological Monographs 27:325–349.

41. Gohl DM, Auch B, Certano A, LeFrançois B, Bouevitch A, Doukhanine E, Fragel C, Macklaim J, Hollister E, Garbe J, Beckman KB. 2021. Dissecting and tuning primer editing by proofreading polymerases. Nucleic Acids Res 49:e87.

42. Lefever S, Pattyn F, Hellemans J, Vandesompele J. 2013. Single-Nucleotide Polymorphisms and Other Mismatches Reduce Performance of Quantitative PCR Assays. Clin Chem 59:1470–1480.

43. Stadhouders R, Pas SD, Anber J, Voermans J, Mes THM, Schutten M. 2010. The Effect of Primer-Template Mismatches on the Detection and Quantification of Nucleic Acids Using the 5′ Nuclease Assay. The Journal of Molecular Diagnostics 12:109–117.

44. Wu J-H, Hong P-Y, Liu W-T. 2009. Quantitative effects of position and type of single mismatch on single base primer extension. Journal of Microbiological Methods 77:267–275.

45. Campos MJ, Quesada A. 2017. Strategies to Improve Efficiency and Specificity of Degenerate Primers in PCR, p. 75–85. In Domingues, L (ed.), PCR: Methods and Protocols. Springer, New York, NY.

46. Nam YR, Lee U, Choi HS, Lee KJ, Kim N, Jang YJ, Joo CH. 2015. Degenerate PCR primer design for the specific identification of rhinovirus C. J Virol Methods 214:15–24.

47. Martin FH, Castro MM, Aboul-ela F, Tinoco I. 1985. Base pairing involving deoxyinosine: implications for probe design. Nucleic Acids Res 13:8927–8938.

48. Watkins NE, SantaLucia J. 2005. Nearest-neighbor thermodynamics of deoxyinosine pairs in DNA duplexes. Nucleic Acids Res 33:6258–6267.

49. Fujiwara H, Fujiwara K, Hashimoto K. 1995. PCR with deoxyinosine-containing primers using DNA polymerases with proofreading activity. Genome Res 4:239–240.

50. Knittel T, Picard D. 1993. PCR with degenerate primers containing deoxyinosine fails with Pfu DNA polymerase. Genome Res 2:346–347.

51. Lozano-Fernandez J. 2022. A Practical Guide to Design and Assess a Phylogenomic Study. Genome Biol Evol 14:evac129.

52. Xu Y, Lewandowski K, Lumley S, Pullan S, Vipond R, Carroll M, Foster D, Matthews PC, Peto T, Crook D. 2018. Detection of Viral Pathogens With Multiplex Nanopore MinION Sequencing: Be Careful With Cross-Talk. Frontiers in Microbiology 9.

53. Aird D, Ross MG, Chen W-S, Danielsson M, Fennell T, Russ C, Jaffe DB, Nusbaum C, Gnirke A. 2011. Analyzing and minimizing PCR amplification bias in Illumina sequencing libraries. Genome Biol 12:R18.

54. Laursen MF, Dalgaard MD, Bahl MI. 2017. Genomic GC-Content Affects the Accuracy of 16S rRNA Gene Sequencing Based Microbial Profiling due to PCR Bias. Front Microbiol 8:1934.

55. Větrovský T, Kolařík M, Žifčáková L, Zelenka T, Baldrian P. 2016. The rpb2 gene represents a viable alternative molecular marker for the analysis of environmental fungal communities. Molecular Ecology Resources 16:388–401.

56. Kebschull JM, Zador AM. 2015. Sources of PCR-induced distortions in high-throughput sequencing data sets. Nucleic Acids Res 43:e143.

57. Shirazi S, Meyer RS, Shapiro B. 2021. Revisiting the effect of PCR replication and sequencing depth on biodiversity metrics in environmental DNA metabarcoding. Ecol Evol 11:15766–15779.

58. Frank JA, Reich CI, Sharma S, Weisbaum JS, Wilson BA, Olsen GJ. 2008. Critical Evaluation of Two Primers Commonly Used for Amplification of Bacterial 16S rRNA Genes. Appl Environ Microbiol 74:2461–2470.

59. Heuer H, Krsek M, Baker P, Smalla K, Wellington EM. 1997. Analysis of actinomycete communities by specific amplification of genes encoding 16S rRNA and gel-electrophoretic separation in denaturing gradients. Appl Environ Microbiol 63:3233–3241.

60. Yu Y, Lee C, Kim J, Hwang S. 2005. Group-specific primer and probe sets to detect methanogenic communities using quantitative real-time polymerase chain reaction. Biotechnology and Bioengineering 89:670–679.

